# TRANCERs: Engineering enhancers into autonomous tissue-specific expression cassettes

**DOI:** 10.1101/2025.10.27.684763

**Authors:** Elisa K. Barrow Molina, Alastair L. Smith, Paco López-Cuevas, Matthew Nicholls, Sumana Sharma, Richard M. White, Thomas A. Milne, Jim R. Hughes

## Abstract

Due to the scarcity of autonomous cell-specific promoters, control of transgene expression remains a major challenge, particularly for therapeutic applications where specificity and compactness are critical. We developed TRANCERs (**TR**anscriptionally **A**utonomous enha**NCER**s), a modular system that converts gene regulatory elements (enhancers) into cell-specific standalone promoters by coupling enhancer activity with known and novel principles of transcript stability and nuclear export. This vastly increases the current repertoire of highly specific promoters available to science. We show TRANCERs drive lineage-restricted expression of genes both *in vitro* and *in vivo*, and that this expression pattern faithfully mirrors the activity of the enhancer element used. Their compact footprint and resistance to epigenetic silencing makes them ideally suited for payload-limited delivery systems such as viral vectors. TRANCERs provide a broadly applicable strategy for precise and programmable control of gene expression, enabling highly specific and customisable expression patterns for both basic research and gene-based therapies.

## Main

Promoters are key regulatory elements that function by stabilising RNA polymerase II (RNAPII) initiation thus promoting transcription across coding regions ^1^. Promoters initiate transcription but alone can rarely generate the cell type- and context-specific expression programs required in multicellular organisms ^2^. This specificity depends on enhancers, non-coding cis-regulatory elements located variably around their target genes, that modulate promoter activity and are themselves frequently transcribed into short, generally bidirectional and unstable enhancer RNAs (eRNAs) ^3–8^. Although the function of eRNAs remains unresolved, their expression strongly correlates with target gene activation and enhancer activity ^9,10^.

Enhancers and promoters share architectural and functional features; however, they differ markedly in their degree of autonomy. The loading of general transcription factors (TFs) on promoters is highly dependent on their interactions with enhancers, which is critical for regulating the level of transcription. However, the loading of TFs on enhancers is inherent in their sequence and is not dependent on other elements and enhancers are more enriched for the binding of cell type-specific factors than promoters ^11,12^. It has also been shown that they can even be moved to completely different genomic environments and still retain their ability to activate with the same cell specificity ^13–16^. The same is also true of the individual enhancer elements that make up super-enhancers. Deletion experiments have shown that the individual elements are active in the absence of the other component enhancers even though such deletions affect the activity of their linked promoter elements ^17,18^. This illustrates that promoters are mostly dependent elements, where enhancers can function as independent regulatory elements.

Importantly, enhancers have been reported to frequently generate highly expressed and cell type specific isoforms of the genes when situated in their introns ^19^. Similarly, an extremely well characterised enhancer from the Alpha Globin gene locus when relocated to a neutral area on mouse Chromosome X was capable of producing high levels of both nascent and stable erythroid-specific transcription and adopted a more promoter-like chromatin signature ^16^. This suggests that enhancers, while intrinsically more autonomous, also have many functional similarities with promoter elements, which can be further induced by cis acting signals in certain genomic contexts. Therefore, if these signals could be understood and co-opted then enhancers could be converted to reliable, autonomous and highly cell type specific sources of transcription. This would be important because while autonomous cell type-specific promoters have been reported, they are rare ^15^. In contrast, there are estimated to be millions of cell type-specific enhancers in the human genome controlling development, differentiation and cellular responses to extrinsic signals ^13^. This therefore holds out the potential to have precise transcriptional control in all conceivable cell types and cells states for which specific enhancers exist. As a unique gene expression profile is the core determinant of cell type or cell state and this expression profile is determined by similarly specific enhancer elements, the suggests that a specific set of enhancers exists for every cell type and cell state in the human body.

Current strategies for achieving cell type-specific expression are highly limited relying on combining characterised enhancers with minimal promoters, but these systems often suffer from leaky expression, unpredictable enhancer–promoter compatibility, repression and limitations on vector size ^20–23^. To address these challenges, we hypothesised that enhancer activity is not inherently weak but rather transcripts get rapidly degraded, and that supplying appropriate downstream processing motifs could convert enhancer transcription into stable, productive output. Here we present TRanscriptionally Autonomous EnhaNCERs (TRANCERs), a platform that transforms enhancer elements into compact, autonomous and cell type/state-specific functional transcriptional sources. We demonstrate that TRANCERs operate robustly across mammalian cell lines and *in vivo*, providing a new framework for programmable and context-specific gene expression.

### Enhancer transcription can be stabilised and made to produce protein coding transcripts via the addition of a range of cis acting sequences

We hypothesised that enhancer transcription is not intrinsically weak, but rather highly unstable, and that this instability could be overcome by incorporating naturally occurring cis-acting sequence elements, such as polyadenylation sites (poly(A)), to protect enhancer-derived transcripts from degradation. However, we anticipated that transcript stabilisation alone would be insufficient to support translation, as efficient engagement with the nuclear export machinery is required for delivery of transcripts to the cytoplasm. We therefore reasoned that both stabilisation signals and mechanisms that actively promote the transcription-export complex (TREX)-mediated nuclear export are necessary to render enhancer-derived transcripts cytoplasmic and translationally competent ^24^. mESCs were initially chosen as a well characterised and tractable system for testing this idea. Active mESC enhancers were identified using TT-seq to sensitively capture nascent, short-lived transcripts as a proxy for enhancer activity and combined with ATAC-seq and ChIP-seq profiling (H3K4me1, H3K27ac, H3K4me3, CTCF) for the annotation of element classes ^25–27^. Focusing on intergenic elements to reduce noise and confounding signals from promoters and gene bodies we yielded a high confidence set of active enhancers which were then compared with the literature to select previously validated enhancer elements (Supplementary Table 1). Enhancer names were adopted from the existing literature. Where there was ambiguous enhancer–gene assignments these were resolved using Micro-Capture-C (MCC), revealing, for example, that a previously annotated *Tmprss13* enhancer instead interacts with *Fxyd6* (Extended Data Fig, 1a). These enhancers were then incorporated into a base TRANCER scaffold containing a synthetic intron for TREX engagement, an mNG (Neon Green) as a reporter, and a polyadenylation signal to stabilise the transcript which was then integrated into a neutral region on chromosome X with no pre-existing function ^16^. All constructs produced clearly detectable protein expression, indicating that enhancer-derived transcripts can acquire promoter-like activity when stabilised with canonical 5′ and 3′ RNA-processing elements. A reversed non-complementary version of the Nanog enhancer was used as a negative control, thus conserving general nucleotide content characteristics, while destroying existing TF binding motifs (Fig. 1a). This negative control showed no protein production and the Cre-LoxP removal of the puromycin cassette (Fig. 1b) did not decrease expression, confirming that the enhancer was the sole transcriptional driver ^28^.

**Fig. 1.**
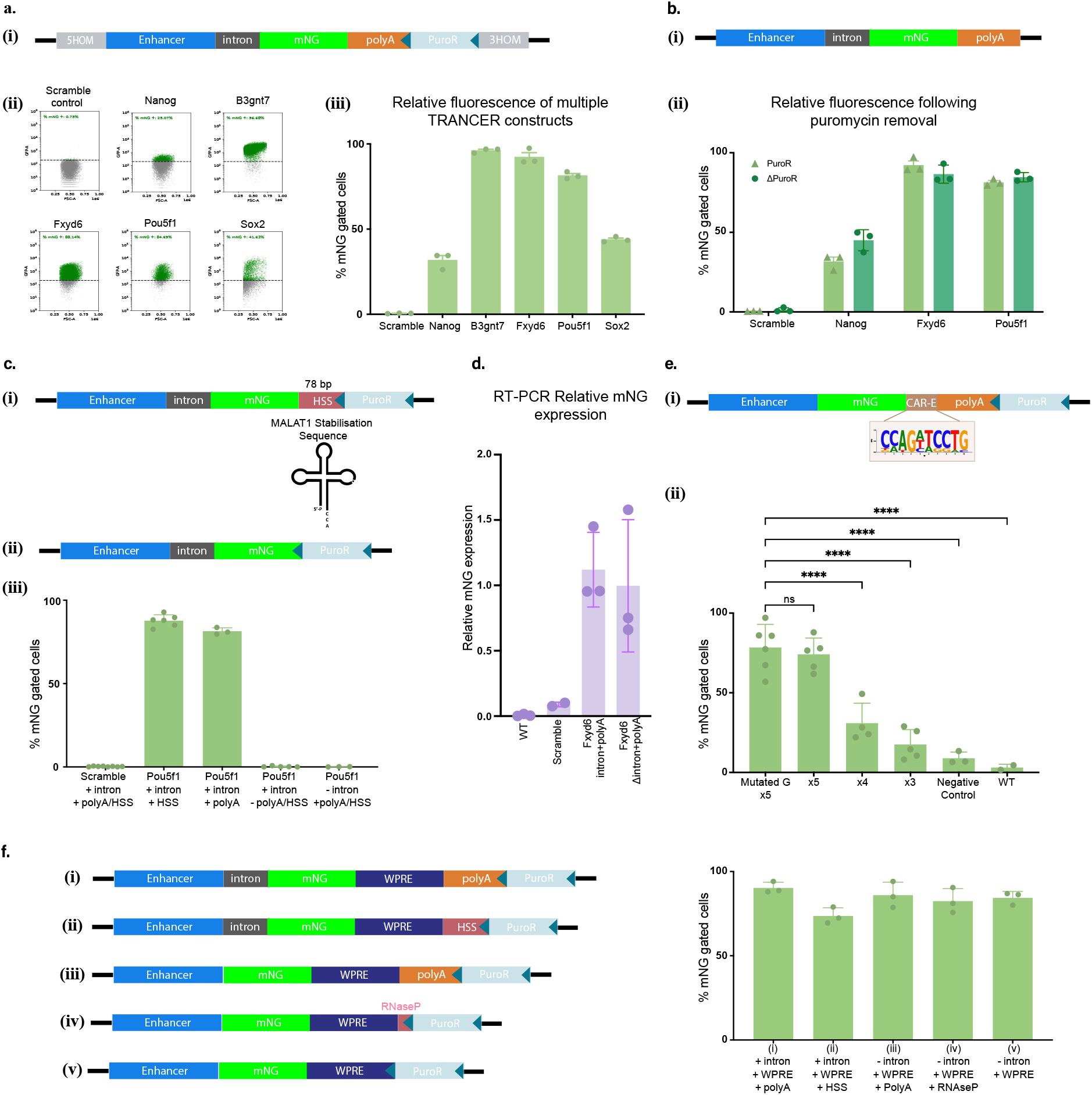
Generation and characterization of TRANCER constructs using multiple approaches. **a, Base TRANCER constructs (intron + polyA)**. (top) Schematic of the TRANCER construct inserted as a single copy into a neutral locus on ChrX. The construct includes a synthetic intron (dark grey box), polyadenylation site (orange box), and an interchangeable enhancer element or control sequence (blue box) that drives specific expression patterns. A floxed puromycin resistance cassette enables selection. Optional 5′ and 3′ homology arms (grey boxes) allow for CRISPR-mediated targeted integration if desired. (i) Flow cytometry (Attune NxT) analysis of Neon Green (mNG) reporter expression, reflecting TRANC-ER-driven protein production. Green and grey populations denote mNG-positive and -negative cells, respectively. Enhancer identity is indicated above each plot. The “Scramble control” corresponds to a Nanog enhancer sequence with disrupted transcription factor binding sites but preserved base composition. (ii) bar graph quantifying mNG-positive cells (%); error bars represent Standard Error of the Mean (SEM). **b, Effect of puromycin cassette excision on mNG expression**. (i) Schematic illustrating removal of the floxed puromycin resistance cassette by Cre recombinase. (ii) Bar plot summarizing flow cytometry data (three biological replicates) comparing TRANCER constructs with (light green triangles) or without (dark green circles) the puromycin cassette. Expression levels remain unchanged, indicating that cassette removal does not affect mNG expression. **c, Replacement of the polyA site with a Hairpin Stabilization Sequence (HSS)**. (i) Schematic showing substitution of the 225 bp polyA signal with a 78 bp HSS derived from the MALAT1 non-coding RNA. (ii) A control construct lacking both polyA and HSS sequences was also generated. (iii) Flow cytometry quantification of mNG fluorescence for each configuration, assessing RNA stabilization efficiency. **d, TRANCER transcript localization and nuclear retention**. qPCR analysis confirming transcript expression and nuclear accumulation without cytoplasmic export, consistent with absent protein production (as in panel c). These data suggest loss of TREX pathway engagement and nuclear retention of the transcript. **e, Rescue of intronless TRANCER expression by CAR-E motifs**. (i)Three to five tandem copies of the 10 bp CAR-E consensus motif were inserted into an intronless TRANCER containing a 3′ polyA site. mNG. (ii) Expression was restored in a copy number–dependent manner, indicating that CAR-E elements can substitute for the intron in promoting expression. **f, Inclusion of Woodchuck Posttranscriptional Regulatory Element (WPRE) restores expression in intron- and polyA-deficient TRANCERs**. (left) Schematic overview of the constructs tested, including: (i) TRANCER with intron and polyA plus WPRE; (ii) HSS-containing construct with WPRE; (iii) construct lacking the intron plus WPRE; (iv) construct lacking polyA but containing an RNAse P cleavage site plus WPRE; and (v) construct lacking both TREX engagement and 3′ stabilization elements in the presence of the WPRE. WPRE inclusion preserves mNG expression even in the absence of intron and polyA sequences. (right) Bar plot quantifying the mNG-positive cells for all constructs.

UTRs are increasingly recognised as key regulators of RNA maturation, stability, localisation, and translation ^29^. To explore alternative 3′ stabilisation strategies, we replaced the poly(A) signal with the conserved MALAT1 triple-helix structure (HSS) ^30^. In addition to protecting against degradation, the HSS has previously been reported to enhance translation, suggesting potential advantages over canonical poly(A) for transgene expression ^31^. While the HSS was equally effective at supporting expression it did not enhance protein output, whereas removal of either stabilisation sequences abolished expression entirely (Fig. 1c).

Eliminating the synthetic intron resulted in nuclear accumulation of transcripts without detectable protein (Fig. 1d, flow cytometry not shown), consistent with the need for TREX mediated, splicing-dependent mRNA export ^32–34^. Inspired by intronless genes that use cytoplasmic accumulation regions (CARs) for export, we replaced the splice site with tandem CAR-E motifs downstream of mNG ^35^. Three to five copies restored protein expression in intronless TRANCERs, with five giving near-complete rescue (Fig. 1e). Unexpectedly, mutation of a conserved CAR-E nucleotide had minimal effect, suggesting that nuclear-to-cytoplasmic export machinery recruitment may rely on structural properties of these repeated motifs rather than strict sequence identity.

Many viruses encode RNA elements that bypass canonical splicing or polyadenylation to enhance heterologous gene expression ^36–40^. One such element, the Woodchuck Hepatitis Virus Post-transcriptional Regulatory Element (WPRE), is widely used in therapeutic vectors to increase transcript stability and protein output ^41–46^. To test it would increase the output of TRANCERs, we incorporated the clinically validated WPREmut6 sequence into constructs containing both an intron and poly(A). Unexpectedly, poly(A) and intron-deficient controls produced mNG at comparable levels to the full constructs showing the WPRE could replace both (Fig. 1f). Thus, the WPRE alone is sufficient to stabilise TRANCER transcripts, promote export, and support translation through a mechanism that remains to be defined.

Here, we have demonstrated that we can stabilise and engage enhancer derived transcription via multiple mechanisms to create productive transcription with a new technology we have called TRANCERs. Overall, the requirements for expression can be split into two main functional requirements: firstly, interaction with the nuclear-to-cytoplasmic export machinery and secondly, 3’ end stabilisation of the transcript.

### Investigating the cell and tissue specific expression of TRANCERs using organoids and zebrafish

Retention of their original tissue-specificity is central to the utility of TRANCERs. We therefore exploited the ability of mESCs’ to differentiate into embryoid bodies (EBs); 3D aggregates that produce a rich mix of cell types from all three germ layers ^47^. During EB differentiation, primitive embryonic erythroid cells emerge, offering a convenient model for rapidly studying enhancer dynamics and specificity ^47,48^. As a benchmark, we deployed the well-characterised α-globin R2 enhancer ^18^. A single R2-driven TRANCER cassette was integrated into the neutral site on chrX (Fig. 2a). Fluorescent reporters were replaced with a truncated human CD19 receptor due to haemoglobin autofluorescence and to enable enrichment of reporter-positive erythroid cells by flow cytometry (Fig. 2b). To profile chromatin accessibility, gene expression, and reporter specificity, we performed a single-cell 10x Multiome assay on EB cells enriched for CD19^+^ cells (∼20% of stained cells). Cell-type annotations (cellxgene) identified expected cells from all germ-layer lineages (Fig. 2c). The UMAP in Fig. 2d(i), coloured by *Hba-α1* expression, marks the annotated erythroid cluster. UMAPs in Fig. 2d(ii-iii) are coloured according to the expression of the CD19 alone Fig. 2d(ii) or the full cassette including the puromycin resistance gene Fig. 2d(iii). Broader signal in Fig. 2d(ii) reflects expression of the linked puromycin cassette driven by an EF1α core promoter, likely boosted in erythroid cells by enhancer–promoter crosstalk. Pseudobulk ATAC-seq from CD19^+^ cells confirmed erythroid-specific chromatin profiles (Extended Data Fig. 2). As expected, only a small fraction of EB-derived cells was CD19 positive, reflecting the limited representation of terminally differentiated RBC within the mixed germ-layer composition of EBs.

**Fig. 2.**
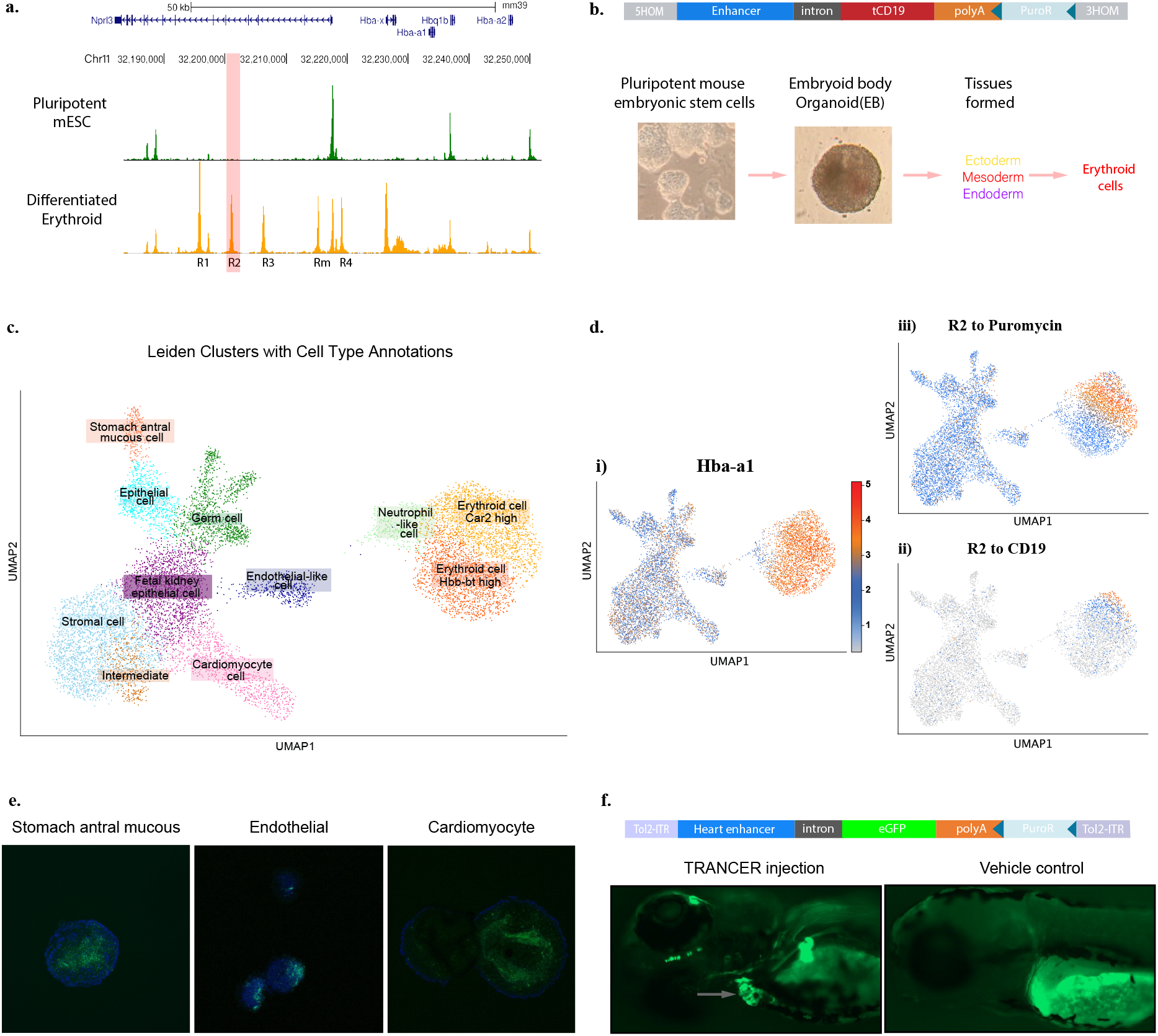
CD19 expression is restricted to erythroid cells in an EB model. **a, ATAC-seq data from differentiated mESC** showing that open chromatin accessibility at the R2 enhancer (highlighted in bold and indicated by the pink bar) is restricted to erythroid cells. **b, Differentiation of mESC into Embryoid Bodies (EBs)** (top) Schematic of a TRANCER construct containing the erythroid-specific R2 enhancer element as the transcriptional driver, with the mNG reporter replaced by the coding sequence of human truncated CD19. (bottom) Stages of erythroid cell generation from mESC in the formation of erythroid cells. Experimental workflow proceeds left to right: formation of embryoid body (EB) organoids, can be followed by immunopurification of red blood cell precursors from the heterogeneous population of ectoderm, mesoderm, and endoderm lineages produced in this system, enabling erythroid enrichment. **c, UMAP projection of full EBs containing the R2–CD19 TRANCER construct**, with annotated cell types corresponding to those identified in wild-type (WT) EBs. **d, UMAP visualizations derived from multiome data demonstrating that the R2–tCD19 TRANCER construct is specifically expressed in erythroid cells**, as evidenced by its colocalization with Hba-a1 expression. (i) Expression of the erythroid marker gene Hba-a1, showing spatial overlap with Cd19-expressing cells. (ii) UMAP of the same dataset, where darker colors represent higher read counts of the inserted R2–tCD19 TRANCER sequence (from the CD19 gene to the puroR cassette). (iii) UMAP representing read counts from CD19 to the polyA tail. **e, Confocal images of fully formed EBs where different TRANCER constructs have been stably integrated into chrX**. (Left) Orthogonal projection image of a representative clone expressing a stomach enhancer. The image shows a diffused yet specific mNG signal in the EB. (Middle) Endothelial enhancer showing expression in three different EBs, localised to one pole of the EB. (Right) Cardiomyocyte enhancer, Single slice from a z-stack displaying mNG in a ring-like formation corresponding to the cardiac tissue region within the EB. **f, Expression of a heart myocardium TRANCER in zebrafish**. Schematic representation of the construct used in this experiment. The TRANCER donor, driven by a heart-specific enhancer, is flanked by Tol2 inverted terminal repeats (ITRs). The reporter used in this construct is eGFP. Fluorescent microscopy images showing tissue-specific eGFP expression localised to the heart myocardium (indicated by grey arrow, left). The right panel shows a negative control embryo that did not receive the injection. Note: mild background autofluorescence is visible in the yolk sac, a common occurrence attributed to the natural composition of yolk lipids and proteins.

Overall, this experiment confirms cell-type specific expression of CD19 in a TRANCER construct driven by the well-characterised R2 enhancer. Additionally, we also selected other enhancers specific to different distinct clusters annotated in our EB multiome analysis and showed distinct expression patterns using confocal imaging (Fig. 2e and Extended Data Fig. 3). To test TRANCER activity in a live vertebrate model, we used zebrafish, which offer exceptional optical access for whole-animal imaging. Candidate myocardium-specific enhancers were identified from public zebrafish scATAC-seq data (10x Multiome) ^49^. The top enhancer candidate was cloned into a base TRANCER (Extended Data Fig.5) and delivered by yolk-sac microinjection using the Tol2 transposase system ^50^. This yielded robust, myocardium-specific TRANCER expression in zebrafish embryos (Fig. 2f), validating *in vivo* enhancer function. Mild ectopic expression near the eye was occasionally observed, likely reflecting mosaic expression in F0 embryos. Beyond the use of natural enhancers, we also have shown that published AI derived-synthetic enhancers can form functional TRANCERs which open up a wide range of possibilities for the generation of highly bespoke expression patterns (Extended Data Fig. 4) ^51^.

### TRANCERs bypass silencing

A major barrier to stable expression of constructs used in gene therapy and synthetic biology is their frequent epigenetic silencing by the Human Silencing Hub (HUSH) complex ^52,53^. To test whether TRANCERs are susceptible to HUSH, we generated base lentiviral TRANCER constructs expressing CD19 under either a synthetic or the erythroid HS2 enhancer element, with a mESC enhancer (*Fxyd6*) as a negative control and transduced them into K562 cells. After puromycin selection for two weeks selection was removed, and cells were monitored weekly by flow cytometry. Whereas traditional constructs often show repression within the first week, TRANCER-expressing cells displayed no loss of CD19 expression over a 12-week period (Fig. 3) ^52,54,55^. Thus, TRANCERs appear resistant to HUSH-mediated silencing and support stable, long-term expression, a major additional advantage over existing transgene technologies.

**Fig. 3.**
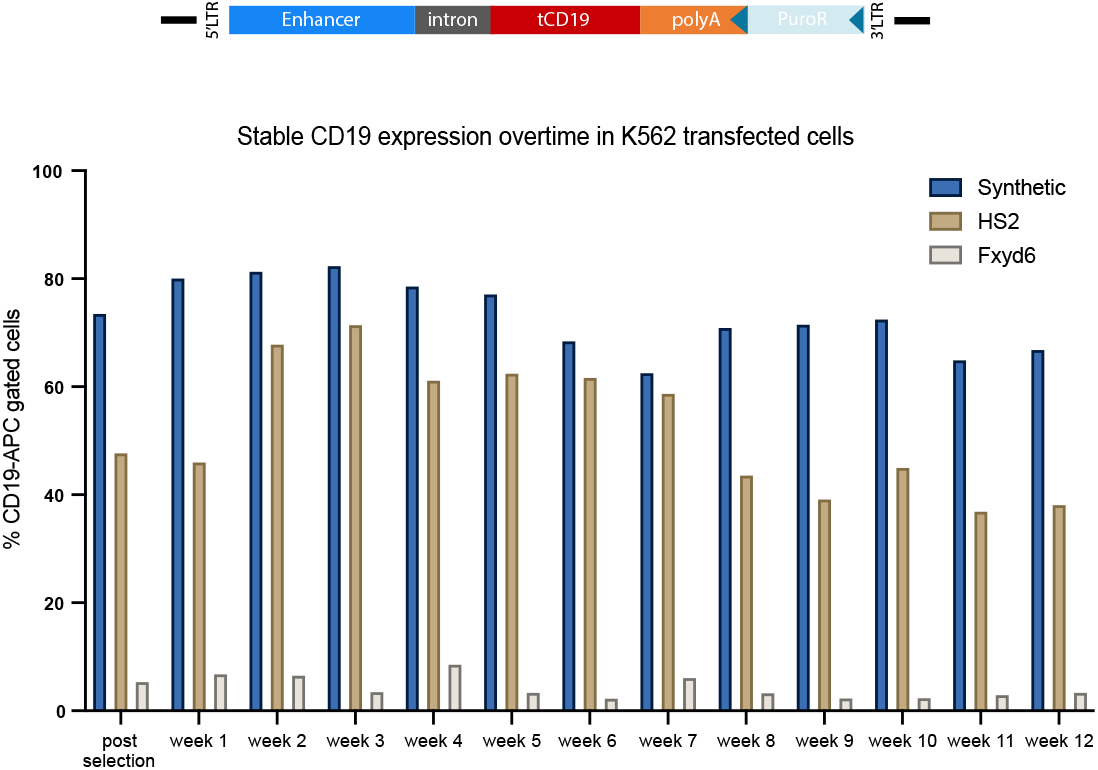
K562 cells were transduced with TRANCER containing either a Synthetic, HS2 or Fxyd6 enhancer driving CD19 expression. Following antibiotic selection for two weeks and complete removal, CD19 surface expression was monitored weekly by flow cytometry, and the percentage of CD19-APC–positive cells was quantified. Both constructs maintained stable expression over several weeks post-selection, with the synthetic enhancer showing consistently higher CD19 expression levels compared to HS2. Data presented as percentage of CD19-positive gated cells over time.

### TRANCERs can be used in many applications: from genome editing to immuno-oncology

The ability to drive protein or RNA expression with high cellular specificity is essential both for therapeutic gene-editing strategies and for basic research. As a platform technology, TRANCERs have the potential to support a broad range of such applications. Here, we illustrate their utility in gene-editing and immuno-oncology contexts.

Many biological processes require the coordinated expression of two transcripts within the same cell particularly when two components must assemble into a functional complex. Enhancers are naturally bidirectional. To determine whether TRANCERs preserve this behaviour, we designed a construct in which a single mESC enhancer (*Fxyd6*) drives expression of two fluorescent proteins in opposite directions with different cis-acting sequences (Fig. 4a). FACS analysis revealed strong co-expression of both fluorophores, confirming efficient bidirectional transcription. A clear use case for such dual expression is CRISPR-mediated editing, which requires both the Cas9 protein and a guide RNA to form an active complex. We tested whether bidirectional TRANCERs could support adenine base editing (ABE), which improves survival and reduces toxicity compared to nuclease-based editing ^56^. Using a GFP reporter system adapted from Katti et al. (2020), we generated an mESC line harbouring a GFP transgene with a premature TAG stop codon at the Rosa26 locus ^57^. GFP expression is abolished in these cells but can be restored if an ABE converts TAG (stop) to TGG (Trp). A bidirectional TRANCER was engineered to express both the ABE and its corresponding sgRNA. Multiple controls were included: no plasmid, a scrambled guide, a previously used scrambled Nanog enhancer (Fig. 1), a cardiomyocyte enhancer (Fig. 2e), and the R2 α-globin enhancer (Fig. 2b). Among these, only the *Fxyd6* mESC enhancer drove expression of both components to repair the premature stop codon and restore GFP fluorescence (Fig. 7b), achieving efficiencies comparable to conventional editing approaches ^57^.

**Fig. 4.**
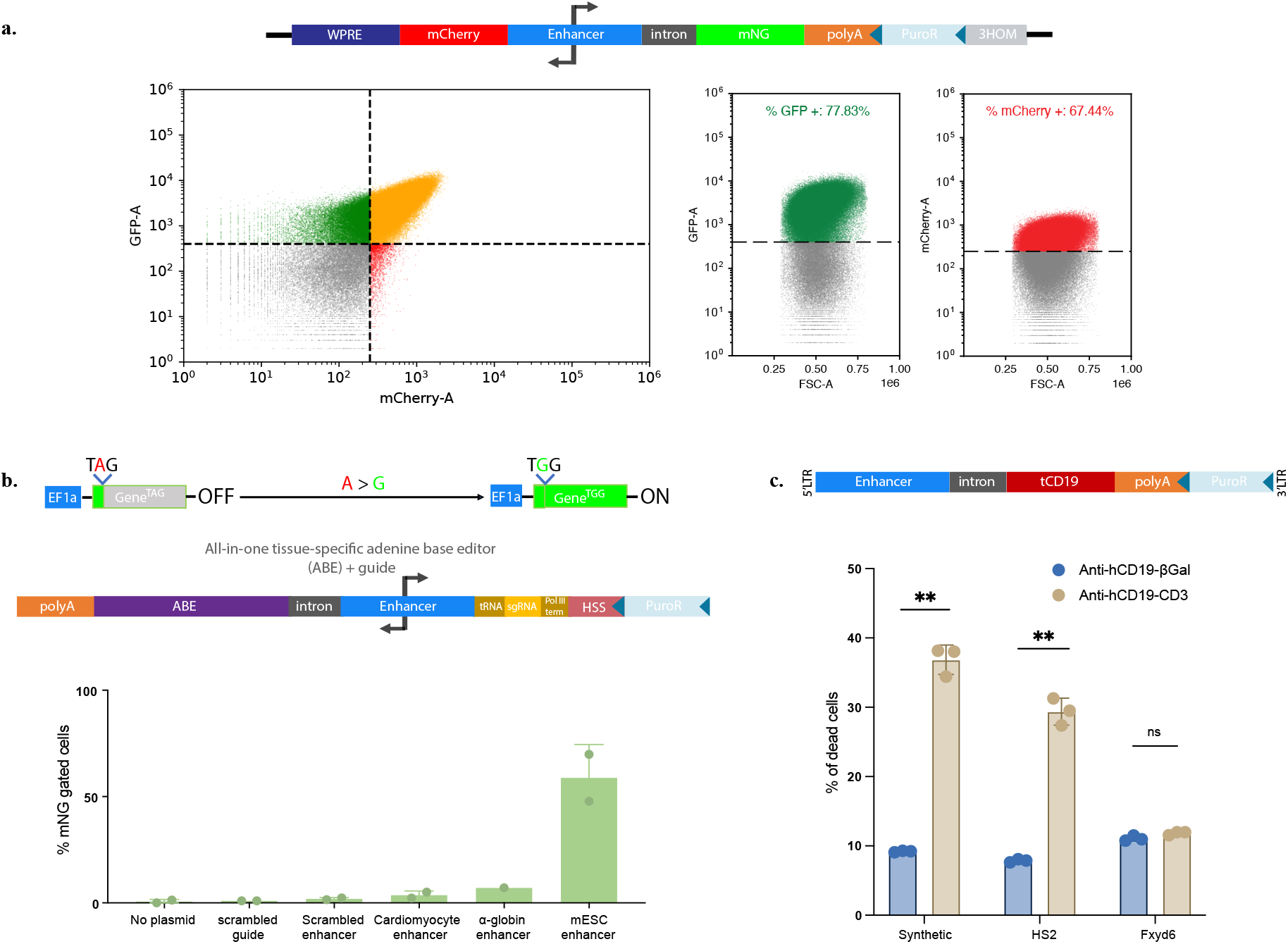
Showing different applications where TRANCER constructs can be used in. **a, Bidirectional TRANCER expressing mCherry and mNG**. (top) Schematic of the TRANCER construct integrated into chrX. From left to right, the construct expresses mCherry, stabilised by the inclusion of a WPRE and driven by the mESC-specific Fxyd6 enhancer. In the opposite direction, mNeonGreen (mNG) expression is stabilised and exported via an intron and a polyadenylation (polyA) sequence. (bottom) Flow cytometry analysis of stably integrated clonal mESCs showing simultaneous bidirectional expression of mCherry and mNG. Double-positive populations correspond to cells co-expressing both fluorophores. Independent single-fluorophore expression profiles are shown to the right. **b, All-in-one bidirectional TRANCER design for tissue-specific base editing**. (top) Schematic of the GFP(TAG) reporter integrated as a single copy at the Rosa26 locus. The premature stop codon (TAG) abolishes GFP translation; base editing that converts A→G restores the wild-type codon (TGG), enabling GFP expression detectable by flow cytometry. (middle) Diagram of the bidirectional lentiviral TRANCER construct used for targeted base editing. One arm expresses a catalytically inactive Cas9 (dCas9)–based editor stabilised by an intron and polyA sequence. The opposite arm expresses the corresponding sgRNA, whose transcription is primarily enhancer-driven and supported by a Pol III full-length tRNA promoter. Background activity is minimised using a Pol III terminator sequence, and transcript stability is enhanced by a 3′ Pol II Hairpin Stabilization Sequence (HSS). A floxed puromycin resistance cassette enables selection. (bottom) Bar plot summarizing flow cytometry data from cells transduced with different TRANCER constructs. From left to right: untransduced control, scrambled sgRNA, scrambled enhancer, cardiomyocyte-specific enhancer (Tbx3, Fig. 3), erythroid enhancer (R2, Fig. 2), and mESC enhancer (Fxyd6). Only the Fxyd6-driven construct restores GFP expression, demonstrating tissue-specific base editing activity. **c, T-cell-mediated killing of target cells expressing CD19 under different TRANCER constructs**. (Top) Schematic depiction of lentiviral TRANCERs driven by either a synthetic, HS2 or Fxyd6 under a base (intron + polyA) approach. Expression of the cell surface marker CD19. (Bottom) Quantification of target-cell death following co-culture with with T cells in the presence of bispecific antibodies (BiTEs). Target cells expressing each enhancer construct were incubated with either control Anti-hCD19–βGal (blue) or Anti-hCD19–CD3 (tan) BiTE. Bars represent mean ± SD of biological replicates. Synthetic and HS2 enhancers support significantly increased killing in the presence of Anti-hCD19–CD3 compared with βGal controls (p < 0.01), whereas the Fxyd6 enhancer shows no significant difference (ns).

As TRANCERs can drive highly selective expression of any cargo, we asked whether they could be used to restore or introduce bispecific antibody (BiTE) target antigens on tumour cells. BiTEs link tumour cells to T cells by binding a tumour antigen and CD3, triggering T-cell activation and cytotoxicity ^58,59^. Using blinatumomab, a CD19×CD3 BiTE used for B-ALL, we tested whether TRANCERs could ectopically express CD19 in CD19-negative malignant cells. Lentiviral TRANCER constructs expressing tCD19 stabilised through the presence of a WPRE were driven by either a synthetic K562 enhancer or HS2 and delivered into K562 cells, which do not normally express CD19. Upon treatment with blinatumomab, but not β-gal control, TRANCER-CD19 expressing cells were selectively eliminated (Fig. 4c). Although effective, BiTEs frequently fail when tumour cells lose the target antigen, leading to relapse ^60,61^. These findings show that TRANCERs can re-sensitise antigen-negative tumour cells to BiTE therapy, supporting their potential in immuno-oncology applications.

## Discussion

We set out to address long-standing limitations in transgene design relevant to both basic research and translational applications, particularly the need for expression systems that maintain high cell-type or state specificity, minimise cargo size, remain unsilenced and are compatible with viral and non-viral delivery strategies. TRANCERs represent a versatile platform capable of meeting these demands.

Enhancer RNAs (eRNAs) have historically been dismissed as unstable by-products of transcriptional noise. However, accumulating evidence demonstrates a tight association between enhancer transcription, enhancer activity, and gene regulation ^9^. Importantly, eRNAs display highly cell-type- and context-specific expression patterns, yet mechanistic insights into their stabilisation remain scarce. Existing evidence for stable eRNAs has largely been observational and has not been translated into transgene applications ^16,19,25,26^.

Here we show that by incorporating stabiliser motifs (e.g. polyA or HSS) and nuclear export elements (e.g. CARs or introns), enhancer sequences can substitute for promoters and support robust, productive transcription. The stabiliser motif protects transcripts from exosome-mediated degradation, while TREX engagement via splicing or cytoplasmic accumulation signals ensures nuclear export and translation. This dual mechanism enables native or synthetic enhancers to generate stable, properly processed RNAs that can be translated into functional protein products, as demonstrated by expression of reporter cargos such as mNG and tCD19. We systematically evaluated combinations of stabilisation and export elements. In mESC enhancers, splice sites engaged TREX while polyadenylation stabilised cargo transcripts. We subsequently discovered that a compact cloverleaf motif (HSS) could substitute for polyadenylation without loss of protein output, and that TREX engagement could be achieved without splicing through tandem repeats of a minimal Cytoplasmic Accumulation Region (CAR). Notably, the essential TRANCER elements can be reduced to ∼128 bp, comparable in size to a minimal CMV promoter yet requiring no additional stabilisation sequence. By contrast, conventional CMV-containing constructs remain dependent on enhancer elements to promote specificity and suffer from mis-expression and silencing. This size advantage could be transformative for delivery of large cargos, particularly where viral packaging capacity is limiting. Similary, we demonstrated that the WPRE element can substitute for both stabilisation and export functions. Although larger (589 bp) than other TRANCER components, WPRE offers a single-insert solution suitable for certain contexts, such as integration into endogenous loci. Future work may enable functional minimisation of this element to further streamline constructs.Additionally, here we show TRANCERs’ stage and tissue specificity conclusively through organoid differentiation of mESCs, as well as *in vivo* using zebrafish (Fig. 2). Finally, the intrinsic bidirectionality of TRANCERs allows for the coordinated expression of multiple transcripts. In summary, TRANCERs provide a modular framework for engineering enhancer-derived transcription into a programmable tool for transgene production. Depending on the application, constructs can be tailored for compactness, cytoplasmic translation, nuclear confinement or the coordinated expression of cargos. This versatility positions TRANCERs as a broadly applicable technology for basic biology, gene therapy, immune-oncology, and genome engineering.

## Materials and methods

### Cell Culture

Mouse embryonic stem cells (E14TG2a.IV; 129/Ola background) were maintained feeder-free on 0.1% gelatin-coated plates in GMEM supplemented with 10% FBS, 1× MEM-NEAA, 2 mM L-glutamine, 1 mM sodium pyruvate, 0.1 mM 2-mercaptoethanol, 1000 U/mL LIF, and 100 U/mL penicillin–streptomycin at 37 °C and 5% CO_2_. Cells were passaged using 0.05% trypsin–EDTA or Accutase and reseeded into fresh medium. K562 human myelogenous leukemia cells were cultured in RPMI-1640 supplemented with 10% FBS and 2 mM L-glutamine and passaged by dilution into pre-warmed medium.

### Primary CD3^+^ T cell culture and T-cell killing assay

Human primary CD3^+^ T cells were isolated from peripheral blood mononuclear cells (PBMCs) using the MojoSort™ Human CD3 Selection Kit (BioLegend, Cat. No. 480133) according to the manufacturer’s instructions. Isolated CD3^+^ T cells were activated and expanded using Dynabeads™ Human T-Activator CD3/CD28 (Life Technologies, Cat. No. 11132D) at a beadto-cell ratio of 1:1 (25 µL beads per 1 × 10^6^ cells) in 24-well tissue culture flasks. Cells were cultured for 5 days, after which activation beads were removed, and the cells were further expanded for an additional 4 days. Expanded CD3^+^ T cells were used for subsequent experiments. All primary T cells were cultured in RPMI 1640 medium (Gibco) supplemented with 10% (v/v) human serum, 1% (v/v) penicillin/streptomycin/neomycin (final concentrations: 50 U/mL penicillin, 50 µg/mL streptomycin, and 100 µg/mL neomycin), 1% (v/v) L-glutamine, 1% (v/v) HEPES, and 0.1% (v/v) β-mercaptoethanol (collectively referred to as “H10 medium”). H10 medium was supplemented with recombinant human IL-2 at 50 U/mL.

Primary CD3^+^ T cells were labelled with CellTrace™ Yellow and co-cultured with engineered K562 target cells expressing CD19 under different enhancer elements at an effector-to-target ratio of 2.5:1. Target cells were incubated with 5 ng/mL of either control anti-hCD19–βGal (bimab-cd19bg-01, InvivoGen) or anti-hCD19–CD3 (bimab-cd19cd3-01, InvivoGen) bispecific (BiTEs) antibodies. Co-cultures were set up in 96-well plates in triplicate and incubated at 37 °C for 24 h. Cell viability was assessed by flow cytometry using Zombie NIR™ live/dead staining, and target cell death was quantified as the percentage of Zombie NIR– positive cells.

### Cloning and Vector Assembly

All constructs were generated using Gibson HiFi assembly according to the manufacturer’s instructions. DNA inserts were either synthesised as gBlocks (IDT) or PCR-amplified from genomic DNA using Q5 High-Fidelity polymerase with gene-specific primers. PCR products and entry vectors were purified prior to assembly. Entry vectors were sourced from in-house stocks (Milne group or MRC WIMM Genome Engineering Core) and prepared either by restriction digestion or PCR amplification. Stable cell lines were generated via CRISPR–Cas9-mediated homology-directed repair (HDR). Guide RNAs and 500 bp homology arms were designed using the IDT Alt-R tool unless otherwise specified. Following assembly, constructs were transformed into *E. coli* DH5α, selected on antibiotic-containing LB agar, and cultured overnight at 37 °C. Plasmid DNA was isolated using standard miniprep procedures, screened by diagnostic digestion, and verified by Sanger sequencing.

### Fluorescence-Activated Cell Sorting (FACS) and Flow Cytometry

FACS was used to identify and isolate successfully transfected cells. Live cells were resuspended in FACS buffer (PBS with 10% FBS) and sorted. Flow cytometry was performed on an Attune NxT flow cytometer to assess TRANCER signal expression and intensity in either 96-well plates or individual tubes. Wild-type, non-expressing cells were used for gating, and cellular debris and doublets were excluded based on forward (FSC) and side scatter (SSC) parameters. Data were analysed using FlowCytometryTools for gating and PRISM software for plotting replicate results.

### Quantitative Reverse Transcription PCR (RT-qPCR)

Total RNA was extracted from cells using a PCR purification kit and reverse-transcribed into cDNA using SuperScript™ III Reverse Transcriptase according to the manufacturers’ instructions. Quantitative PCR was performed using gene-specific primers and a SYBR Green– based master mix with Low ROX normalization on a QuantStudio™ Real-Time PCR System. Reactions (20 µL) contained 15 µL master mix with primers at 500 nM and 5 µL cDNA and were run in technical triplicates. Thermal cycling conditions consisted of 95 °C for 10 min followed by 40 cycles of 95 °C for 15 s and 60 °C for 1 min. Melt curve analysis was used to confirm amplification specificity. Primer efficiencies were assessed using 3-fold serial dilutions of reference cDNA. Relative gene expression was calculated using the ΔΔCt method with normalization to housekeeping genes.

### Differentiation of mESCs into Embryoid Bodies (EBs)

Embryoid body (EB) cultures were generated as previously described Francis *et al*., (2022) ^48^. Forty-eight hours prior to differentiation, mESCs were passaged into Iscove’s Modified Dulbecco’s Medium (IMDM) supplemented with LIF. To initiate differentiation (day 0), cells were dissociated with trypsin and resuspended in freshly prepared differentiation medium consisting of 84.5 mL IMDM, 16.5 mL heat-inactivated ΔFBS, 5.5 mL PFHM II, 1 mL GlutaMAX, 1 mL transferrin, 1 mL amino acids, and 0.0026% (v/v) monothioglycerol (MTG). Single-cell suspensions were plated in 10 cm non-adherent dishes at 3×10^3^ cells per dish and cultured for seven days with daily gentle agitation to prevent EB attachment. EBs were harvested and dissociated in 0.25% trypsin for 5 min at 37 °C, and resulting single cells were analysed by FACS to detect mNG^+^ populations.

### Confocal imaging of Embryoid Bodies (EBs)

Samples were fixed with 4% PFA for 20 min on ice and mounted in VECTASHIELD HardSet Antifade Mounting Medium with DAPI (H-1500, Vector Laboratories) on SuperFrost slides (Thermo Scientific SuperFrost Plus, Thermo Scientific) that were modified using autoclave tape to include a shallow well at the centre. Samples were dehydrated using 100% methanol and were cleared using BABB (1:2 benzyl alcohol:benzyl benzoate) before the coverslip was added and sealed with clear nail polish. Images were acquired at room temperature using a Zeiss LSM 900 upright microscope with a 25x oil immersion objective (LD LCI PA 25x/0.8 DIC) and were reconstructed as maximum intensity projections using ZEN 3.1 (Blue Edition) software.

### Lentivirus Production and Transduction

Lentiviral particles were produced in HEK293T cells using a three-plasmid second-generation packaging system. The packaging plasmid psPAX2 (Addgene #12260) or integrase-deficient psPAX2-D64V (Addgene #63586), the envelope plasmid pMD2.G (Addgene #12259), and the transfer vector Lenti MCP-LSD1 Hygro (Addgene #138457) were co-transfected using Lipofectamine 2000. Viral supernatants were collected 48 and 72 h post-transfection, filtered (0.45 μm), flash-frozen, and stored at −80 °C. Insert sequences were cloned into the transfer vector via NheI/KpnI digestion and Gibson HiFi assembly, and constructs were verified by Sanger sequencing. For transduction of suspension cells, spinfection was performed at 1000 × g for 90 min at 32 °C in the presence of 8 μg/mL polybrene, followed by antibiotic selection (puromycin or blasticidin) 24 h later. For mESC transduction, Retronectin-coated plates were used to enhance infection. Viral particles were bound to plates by centrifugation at 2000 × g for 2 h at 32 °C, after which 1×10^5^ target cells were added and spinfected at 1000 × g for 1 h. Cells were subsequently cultured at 37 °C for recovery and downstream assays.

### Zebrafish husbandry, ethics, and transgenesis

Adult zebrafish (*Danio rerio*) were maintained under standard laboratory. All experiments were reviewed and approved by the University of Oxford Animal Welfare and Ethical Review Body (AWERB) and conducted in accordance with the UK Animals (Scientific Procedures) Act 1986 under Home Office Project Licence PPL PP2565788. The wild-type strain used in this study was AB/Tübingen (AB/TU).

For transgenesis, 1 nL of injection mix containing 25 pg plasmid DNA and 20 pg Tol2 transposase mRNA was injected into the yolk sac of one-cell stage embryos as previously described ^62^. Injected larvae were anaesthetized at 4 days post-fertilization with 0.16 mg ml^-1^ tricaine methanesulfonate (MS-222; Sigma) and imaged using a Zeiss Axio Zoom.V16 microscope with Zen 3.11 software.

### Single-Cell Multiome (ATAC + RNA) Sequencing

mESCs were differentiated into embryoid bodies (EBs) as described above. FACS was performed on wild-type and R2-CD19 lines, collecting ∼1×10^5^ cells per sample. CD19^+^ cells were enriched using APC anti-human CD19 antibody (BioLegend, #302211). Nuclei isolation was performed according to the 10x Genomics Chromium Single Cell Multiome ATAC + Gene Expression protocol (CG000365, Rev C; low-cell input workflow). Briefly, cells were washed and pelleted (300 × g, 5 min, 4 °C), resuspended in PBS + 0.04% BSA, and lysed in chilled Lysis Buffer. Nuclei were subsequently washed and resuspended in Diluted Nuclei Buffer as per the manufacturer’s instructions. The resulting nuclei suspensions were immediately used for 10x Genomics library preparation and sequencing.

### Bioinformatic Analysis

All sequencing datasets were processed using the *Seqnado* pipeline (https://pypi.org/project/seqnado) with default parameters and aligned to either the mm39 or hg38 genome builds. **ChIP-seq, ChIPmentation, and ATAC-seq** reads were trimmed with *Trim Galore*, aligned using *Bowtie2*, and visualised as normalised bigwig files via *deepTools* on the UCSC Genome Browser. Peaks were called using *LanceOtron*. **TT-seq** data was aligned with *STAR*, quantified using *featureCounts*, and visualised with *deepTools* for strand-specific transcription profiles. Micro-Capture-C data were aligned with Bowtie2 and processed using the Seqnado MCC workflow. Capture viewpoints were defined using a probe annotation file and reads overlapping capture oligonucleotides were assigned to grouped viewpoints. Interaction profiles were generated for each viewpoint and exported as genome browser–compatible tracks for visualization on the UCSC Genome Browser.10x **Single-cell Multiome (ATAC + RNA)** data were processed with *Cell Ranger ARC* using a custom mm39 reference including the R2-tCD19 locus. Downstream analysis was performed using *muon, ArchR*, and *Seurat*. Quality control included ambient RNA removal (*CellBender*), doublet exclusion (*Scrublet* and *ArchR*), and filtering for cells with >1000 detected genes. Dimensionality reduction and integration of RNA and ATAC modalities were performed with UMAP and *muon*, and cell types were annotated using *cellxgene* transfer learning models.

## Acknowledgments

The authors would like to express their gratitude to all colleagues who contributed to this work, in particular the flow cytometry facility at the Medical Research Council (MRC) Weatherall Institute of Molecular Medicine (WIMM) for providing cell analysis services. This work was supported by a Wellcome Strategic Award (106130/Z/14/Z to J.R.H.) a Wellcome Discovery Award (225220/Z/22/Z), and the Medical Research Council (MRC) (MC_UU_00016/14 and MC_UU_00029/3 to J.R.H., MC_UU_00016/6 to T.A.M and A.L.S). E.K.B.M was supported by the Biotechnology and Biological Sciences Research Council and the Reuben Foundation. M.N was supported by the NC3Rs Training Grant NC/X00158X/1. R.M.W is funded through the Ludwig Institute for Cancer Research and the Melanoma Research Alliance. PLC is funded through the Ludwig Institute for Cancer Research (LICR) and the Oxford Centre for Cancer Early Detection and Prevention (OxCODE). S.S is funded by the Wellcome Trust (317519/Z/24/Z).

## Contributions

E.K.B.M. and J.R.H. conceived the study and designed the experiments. T.A.M. and A.L.S contributed to experimental design. E.K.B.M. performed experiments and data analysis. A.L.S. provided data analysis. P.L.C. performed the zebrafish experiments. M.N., in collaboration with E.K.B.M., performed confocal imaging experiments. S.S. provided key reagents. R.M.W. provided financial support for the zebrafish experiments performed by P.L.C. The manuscript was written by E.K.B.M., J.R.H. and T.A.M. All authors reviewed and approved the manuscript. J.R.H. and T.A.M. supervised the research.

## Competing interests

J.R.H. is founder, shareholder, director, and paid consultant of Nucleome Therapeutics. J.R.H hold patents for Capture-C (WO2017068379A1, EP3365464B1, US10934578B2). T.A.M. is a paid consultant for, and shareholder in, Dark Blue Therapeutics Ltd. The remaining authors declare no competing financial interests. The remaining authors declare no competing interests. A patent has been applied for under international application number (PCT/GB2025/052425) by the University of Oxford.

## Data availability

All sequencing data generated in this study have been deposited in the Gene Expression Omnibus (GEO). ATAC-seq data from wild-type mESCs are available under accession GSE314141. ChIPmentation data for H3K27ac in wild-type mESCs are available under GSE314143. ChIP–seq data for CTCF in wild-type mESCs are available under GSE314370. TT-seq data from wild-type mESCs are available under GSE314372. MCC data are available under GSE314396. Multiome data from wild-type and R2-tCD19 TRANCER mESCs are available under GSE315186. Reviewer access credentials are provided upon request and during the peer-review process.

## Notes

### Summary of Updates

Changed the manuscript to a newer, more completed version. Changed abstract. Changed competing interest as it was missing the patent number for this work.

## References

1. Haberle, V. & Stark, A. Eukaryotic core promoters and the functional basis of transcription initiation. Nat. Rev. Mol. Cell Biol. 19, 621–637 (2018).

2. Cooper, S. J., Trinklein, N. D., Anton, E. D., Nguyen, L. & Myers, R. M. Comprehensive analysis of transcriptional promoter structure and function in 1% of the human genome. Genome Res. 16, 1 (2006).

3. Pennacchio, L. A., Bickmore, W., Dean, A., Nobrega, M. A. & Bejerano, G. Enhancers: five essential questions. Nat. Rev. Genet. 14, 288 (2013).

4. Lettice, L. A. et al. A long-range Shh enhancer regulates expression in the developing limb and fin and is associated with preaxial polydactyly. Hum. Mol. Genet. 12, 1725– 35 (2003).

5. de Santa, F. et al. A large fraction of extragenic RNA Pol II transcription sites overlap enhancers. PLoS Biol. 8, (2010).

6. Kwak, H., Fuda, N. J., Core, L. J. & Lis, J. T. Precise maps of RNA polymerase reveal how promoters direct initiation and pausing. Science (1979). 339, 950–953 (2013).

7. Andersson, R. et al. Nuclear stability and transcriptional directionality separate functionally distinct RNA species. Nat. Commun. 5, (2014).

8. Arnold, P. R., Wells, A. D. & Li, X. C. Diversity and Emerging Roles of Enhancer RNA in Regulation of Gene Expression and Cell Fate. Front. Cell Dev. Biol. 7, 503820 (2020).

9. Vernimmen, D., Gobbi, M.De, Sloane-Stanley, J. A., Wood, W. G. & Higgs, D. R. Long-range chromosomal interactions regulate the timing of the transition between poised and active gene expression. EMBO J. 26, 2041 (2007).

10. Plant, K. E., Routledge, S. J. E. & Proudfoot, N. J. Intergenic transcription in the human beta-globin gene cluster. Mol. Cell. Biol. 21, 6507–6514 (2001).

11. Arnold, C. D. et al. Genome-wide quantitative enhancer activity maps identified by STARR-seq. Science 339, 1074–1077 (2013).

12. Long, H. K., Prescott, S. L. & Wysocka, J. Ever-Changing Landscapes: Transcriptional Enhancers in Development and Evolution. Cell 167, 1170–1187 (2016).

13. Van Arensbergen, J. et al. Genome-wide mapping of autonomous promoter activity in human cells. Nat. Biotechnol. 35, 145 (2016).

14. Arnold, M. & Stengel, K. R. Emerging insights into enhancer biology and function. Transcription 14, 68 (2023).

15. Heinz, S., Romanoski, C. E., Benner, C. & Glass, C. K. The selection and function of cell type-specific enhancers. Nat. Rev. Mol. Cell Biol. 16, 144 (2015).

16. Georgiades, E. et al. Active regulatory elements recruit cohesin to establish cell specific chromatin domains. Scientific Reports 2025 15:1 15, 1–16 (2025).

17. Hay, D. et al. Genetic dissection of the α-globin super-enhancer in vivo. Nat. Genet. 48, 895–903 (2016).

18. Blayney, J. W. et al. Super-enhancers include classical enhancers and facilitators to fully activate gene expression. Cell 186, 5826-5839.e18 (2023).

19. Kowalczyk, M. S. et al. Intragenic enhancers act as alternative promoters. Mol. Cell 45, 447–458 (2012).

20. Bhatt, B., García-Díaz, P. & Foight, G. W. Synthetic transcription factor engineering for cell and gene therapy. Trends Biotechnol. 42, 449–463 (2024).

21. Hernandez-Garcia, C. M. & Finer, J. J. Identification and validation of promoters and cis-acting regulatory elements. Plant Science 217-218, 109–119 (2014).

22. Milone, M. C. & O’Doherty, U. Clinical use of lentiviral vectors. Leukemia 2018 32:7 32, 1529–1541 (2018).

23. Kim, H., Kim, M., Im, S. K. & Fang, S. Mouse Cre-LoxP system: general principles to determine tissue-specific roles of target genes. Lab. Anim. Res. 34, 147–159 (2018).

24. Pacheco-Fiallos, B. et al. mRNA recognition and packaging by the human transcription-export complex. Nature 616, 828–835 (2023).

25. Zhang, Z. et al. Transcriptional landscape and clinical utility of enhancer RNAs for eRNA-targeted therapy in cancer. Nature Communications 2019 10:1 10, 1–12 (2019).

26. Fukaya, T. Enhancer dynamics: Unraveling the mechanism of transcriptional bursting. Sci. Adv. 9, (2023).

27. Song, C. et al. eRNAbase: a comprehensive database for decoding the regulatory eRNAs in human and mouse. Nucleic Acids Res. 52, D81–D91 (2024).

28. Sauer, B. Functional expression of the cre-lox site-specific recombination system in the yeast Saccharomyces cerevisiae. Mol. Cell. Biol. 7, 2087 (1987).

29. Zhao, W., Blagev, D., Pollack, J. L. & Erle, D. J. Toward a Systematic Understanding of mRNA 3′ Untranslated Regions. Proc. Am. Thorac. Soc. 8, 163 (2011).

30. Brown, J. A., Valenstein, M. L., Yario, T. A., Tycowski, K. T. & Steitz, J. A. Formation of triple-helical structures by the 3′-end sequences of MALAT1 and MENβ noncoding RNAs. Proc. Natl. Acad. Sci. U. S. A. 109, 19202–19207 (2012).

31. Wilusz, J. E. et al. A triple helix stabilizes the 3′ ends of long noncoding RNAs that lack poly(A) tails. Genes Dev. 26, 2392 (2012).

32. Brown, T. A. Synthesis and Processing of RNA. https://www.ncbi.nlm.nih.gov/books/NBK21132/ (2002).

33. Okamura, M., Inose, H. & Masuda, S. RNA Export through the NPC in Eukaryotes. Genes 2015, Vol. 6, Pages 124-149 6, 124–149 (2015).

34. Khan, M., Hou, S., Chen, M. & Lei, H. Mechanisms of RNA export and nuclear retention. Wiley Interdiscip. Rev. RNA 14, e1755 (2023).

35. Lei, H., Zhai, B., Yin, S., Gygi, S. & Reed, R. Evidence that a consensus element found in naturally intronless mRNAs promotes mRNA export. Nucleic Acids Res. 41, 2517 (2013).

36. Guang, S. & Mertz, J. E. Pre-mRNA processing enhancer (PPE) elements from intronless genes play additional roles in mRNA biogenesis than do ones from intron-containing genes. Nucleic Acids Res. 33, 2215–2226 (2005).

37. Mouzannar, K., Schauer, A. & Liang, T. J. The Post-Transcriptional Regulatory Element of Hepatitis B Virus: From Discovery to Therapy. Viruses 2024, Vol. 16, Page 528 16, 528 (2024).

38. Rausch, J. W. & le Grice, S. F. J. HIV Rev Assembly on the Rev Response Element (RRE): A Structural Perspective. Viruses 2015, Vol. 7, Pages 3053-3075 7, 3053–3075 (2015).

39. Pasquinelli, A. E. et al. The constitutive transport element (CTE) of Mason-Pfizer monkey virus (MPMV) accesses a cellular mRNA export pathway. EMBO J. 16, 7500 (1997).

40. Gales, J. P., Kubina, J., Geldreich, A. & Dimitrova, M. Strength in Diversity: Nuclear Export of Viral RNAs. Viruses 12, (2020).

41. Hope, T. J., Zufferey, R., TronoJohn, D. & Donello, E. RNA export element. (1997).

42. Mastroyiannopoulos, N. P., Feldman, M. L., Uney, J. B., Mahadevan, M. S. & Phylactou, L. A. Woodchuck post-transcriptional element induces nuclear export of myotonic dystrophy 3′ untranslated region transcripts. EMBO Rep. 6, 458 (2005).

43. Schambach, A. et al. Woodchuck hepatitis virus post-transcriptional regulatory element deleted from X protein and promoter sequences enhances retroviral vector titer and expression. Gene Therapy 2006 13:7 13, 641–645 (2005).

44. Zufferey, R., Donello, J. E., Trono, D. & Hope, T. J. Woodchuck Hepatitis Virus Posttranscriptional Regulatory Element Enhances Expression of Transgenes Delivered by Retroviral Vectors. J. Virol. 73, 2886 (1999).

45. Brun, S., Faucon-Biguet, N. & Mallet, J. Optimization of transgene expression at the posttranscriptional level in neural cells: implications for gene therapy. Molecular Therapy 7, 782–789 (2003).

46. Ye, L. et al. Enhancement of plasmid-mediated stable gene expression by woodchuck hepatitis virus post-transcriptional regulatory element (WPRE) in human embryonic kidney (HEK293) cells. Afr. J. Biotechnol. 11, 10792–10798 (2012).

47. Oudelaar, A. M., Beagrie, R. A., Kassouf, M. T. & Higgs, D. R. The mouse alpha-globin cluster: a paradigm for studying genome regulation and organization. Curr. Opin. Genet. Dev. 67, 18–24 (2021).

48. Francis, H. S. et al. Scalable in vitro production of defined mouse erythroblasts. PLoS One 17, (2022).

49. Lange, M. et al. A multimodal zebrafish developmental atlas reveals the state-transition dynamics of late-vertebrate pluripotent axial progenitors. Cell 187, 6742-6759.e17 (2024).

50. Urasaki, A., Morvan, G. & Kawakami, K. Functional Dissection of the Tol2 Transposable Element Identified the Minimal cis-Sequence and a Highly Repetitive Sequence in the Subterminal Region Essential for Transposition. Genetics 174, 639 (2006).

51. Gosai, S. J. et al. Machine-guided design of cell-type-targeting cis-regulatory elements. Nature 2024 634:8036 634, 1211–1220 (2024).

52. Spencley, A. L. et al. Co-transcriptional genome surveillance by HUSH is coupled to termination machinery. Mol. Cell 83, 1623-1639.e8 (2023).

53. Müller, I. & Helin, K. Keep quiet: the HUSH complex in transcriptional silencing and disease. Nature Structural & Molecular Biology 2024 31:1 31, 11–22 (2024).

54. Moore, T. V. et al. HDAC inhibition prevents transgene expression downregulation and loss-of-function in T-cell-receptor-transduced T cells. Mol. Ther. Oncolytics 20, 352 (2021).

55. Cabrera, A. et al. The sound of silence: transgene silencing in mammalian cell engineering. Cell Syst. 13, 950 (2022).

56. Gaudelli, N. M. et al. Programmable base editing of A•T to G•C in genomic DNA without DNA cleavage. Nature 2017 551:7681 551, 464–471 (2017).

57. Katti, A. et al. GO: a functional reporter system to identify and enrich base editing activity. Nucleic Acids Res. 48, 2841–2852 (2020).

58. Huehls, A. M., Coupet, T. A. & Sentman, C. L. Bispecific T cell engagers for cancer immunotherapy. Immunol. Cell Biol. 93, 290 (2014).

59. Einsele, H. et al. The BiTE (bispecific T-cell engager) platform: Development and future potential of a targeted immuno-oncology therapy across tumor types. Cancer 126, 3192–3201 (2020).

60. Wang, J. et al. Clinical development of immuno-oncology therapeutics. Cancer Lett. 617, 217616 (2025).

61. Trabolsi, A., Arumov, A. & Schatz, J. H. T Cell–Activating Bispecific Antibodies in Cancer Therapy. The Journal of Immunology 203, 585–592 (2019).

62. Xin, Y. & Duan, C. Microinjection of Antisense Morpholinos, CRISPR/Cas9 RNP, and RNA/DNA into Zebrafish Embryos. Methods Mol. Biol. 1742, 205–211 (2018).

