## Supplemental files for "TRANCERs: Engineering enhancers into autonomous tissue-specific expression cassettes"

**Extended Data Table 1 | Selected mESC enhancers.** *Tmprss13* was renamed to *Fxyd6*.

| Gene name | Distance from TSS | Gene coordinates (mm39) | Size (bp) |
|---|---|---|---|
| *Nanog* | 4560 | chr6:122,679,374-122,679,872 | 499 |
| *B3gnt7* | 3123 | chr1:86,238,385-86,239,098 | 714 |
| *Tmprss13** | 1337 | chr9:45,260,481-45,261,718 | 1238 |
| *Pou5f1* | 2007 | chr17:35,814,791-35,815,159 | 368 |
| *Sox2* | 108618 | chr3:34,799,284-34,799,705 | 421 |

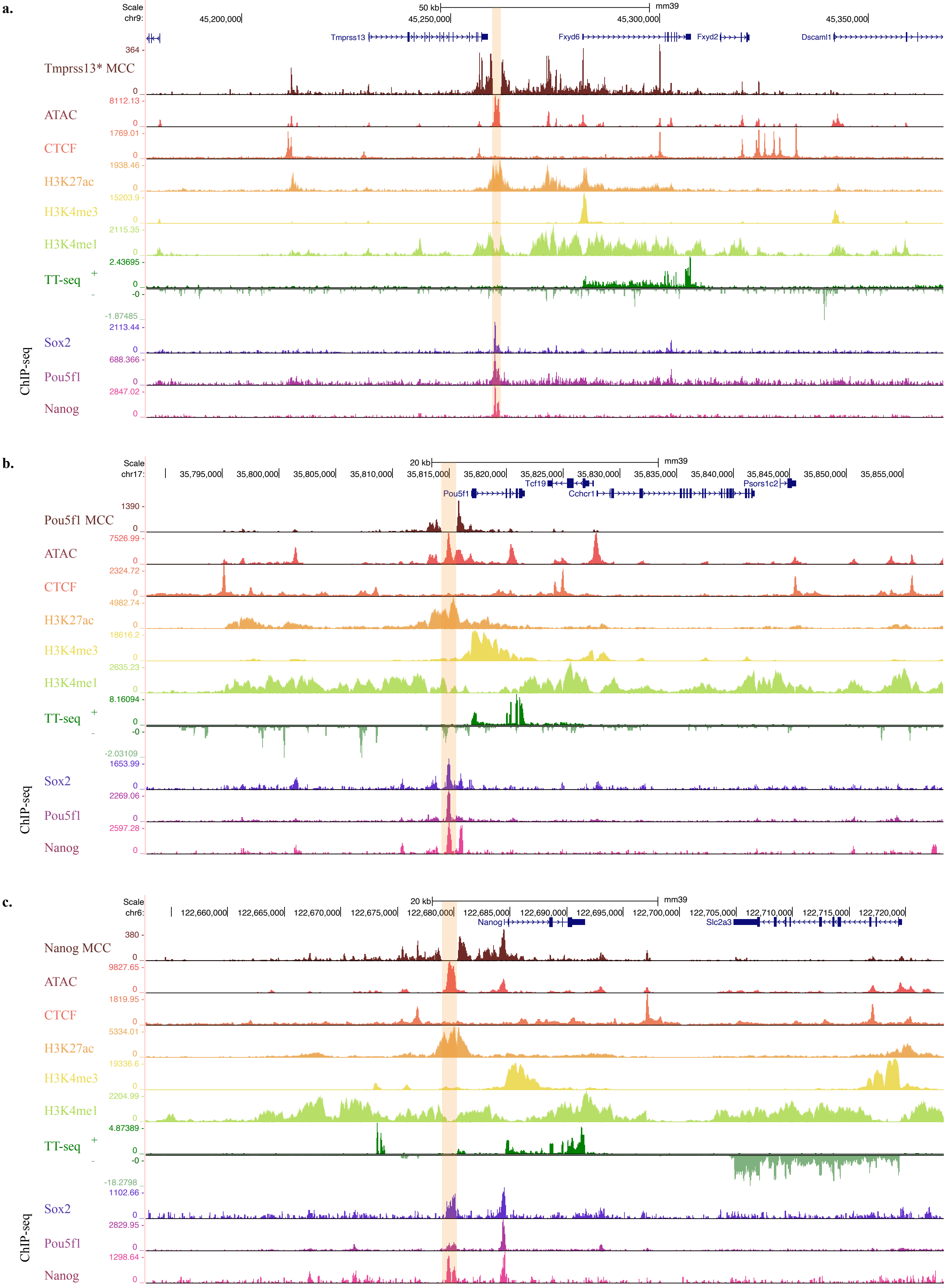
a.
Scale
chr9:
50 kb
mm39
45,200,000
45,250,000
45,300,000
45,350,000
Tmprss13
Fxyd6
Fxyd2
Dscaml1
Tmprss13* MCC
ATAC
CTCF
H3K27ac
H3K4me3
H3K4me1
TT-seq
Sox2
Pou5f1
Nanog
ChIP-seq
b.
Scale
chr17:
20 kb
mm39
35,795,000
35,800,000
35,805,000
35,810,000
35,815,000
35,820,000
35,825,000
35,830,000
35,835,000
35,840,000
35,845,000
35,850,000
35,855,000
Pou5f1
Tcf19
Cchcr1
Psors1c2
Pou5f1 MCC
ATAC
CTCF
H3K27ac
H3K4me3
H3K4me1
TT-seq
Sox2
Pou5f1
Nanog
ChIP-seq
c.
Scale
chr6:
20 kb
mm39
122,660,000
122,665,000
122,670,000
122,675,000
122,680,000
122,685,000
122,690,000
122,695,000
122,700,000
122,705,000
122,710,000
122,715,000
122,720,000
Nanog
Slc2a3
Nanog MCC
ATAC
CTCF
H3K27ac
H3K4me3
H3K4me1
TT-seq
Sox2
Pou5f1
Nanog
ChIP-seq

**d.**

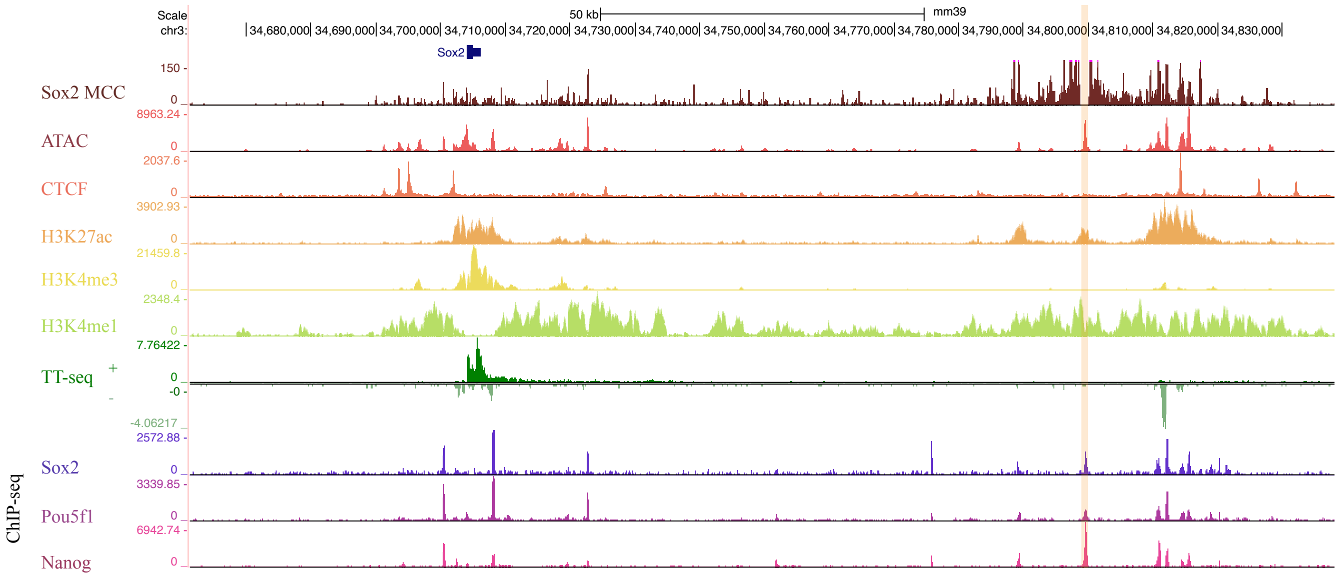


**e.**

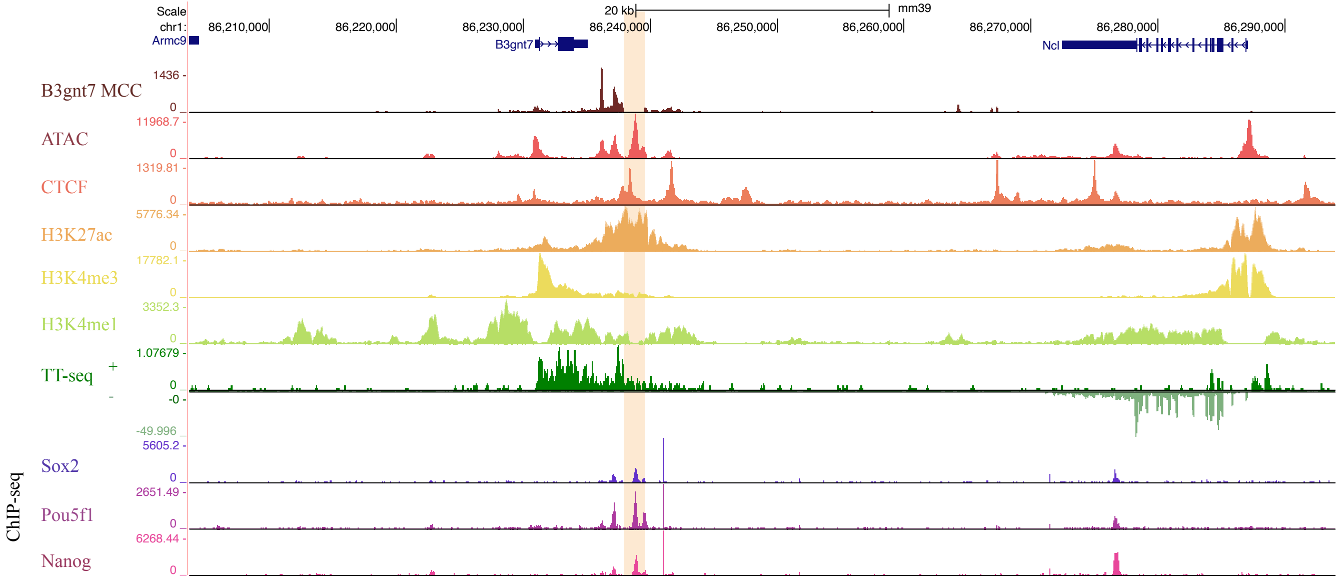


**Extended Data Fig. 1 | UCSC genome tracks displaying (top to bottom) MCC tracks with probes designed to the different enhancers, ATAC-seq, CTCF ChIP-seq, H3K27ac ChIPmentation, H3K4me3 ChIP-seq and H3K4me1 ChIP-seq, TT-seq and ChIP for TF binding of Sox2, Pou5f1 and Nanog.** TT-seq was used to detect unstable eRNAs, Chromatin marks were used in the identification of the different enhancers. To confirm the correct gene assignment of the enhancers (labelled by the highlight) MCC was performed. The *Tmprss13* enhancer was incorrectly assigned in the literature. The *Tmprss13* gene is not active in mESC and the enhancer is instead can be seen to be interacting with the *Fxyd6* gene promoter in the MCC data (a). The additional peaks shown in the MCC tracks align with CTCF peaks and likely represent a subTAD within this region. The *Pou5f1* (b), *Nanog* (c), *Sox2* (d), and *B3gnt7* (e) enhancers all show clear interactions with their assigned genes. H3K4me1 and H3K4me3 ChIP-seq data were obtained from GSE12037, and Sox2, Pou5f1, and Nanog ChIP-seq data were obtained from GSE44286.

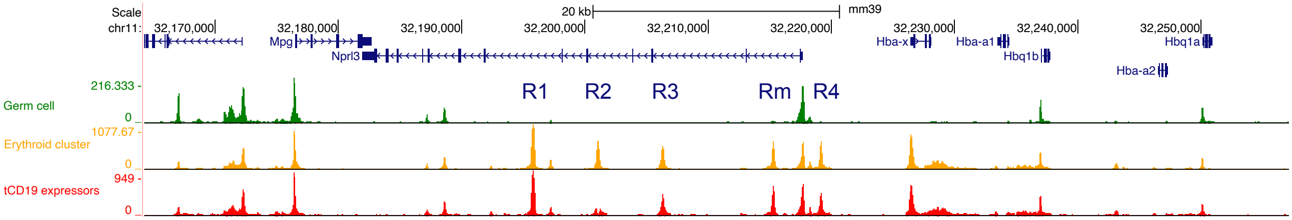


**Extended data Fig. 2 | Pseudobulk tracks derived from WT germ cell and erythroid clusters compared with the CD19 (Fig. 2 panel d (iii) data from TRANCER-containing EBs.** The ATAC-seq accessibility signature is identical between the erythroid cluster and CD19-expressing cells, confirming erythroid-specific activation of the R2 TRANCER. Note that multimapping issues resulted in R2 reads being excluded from the track.

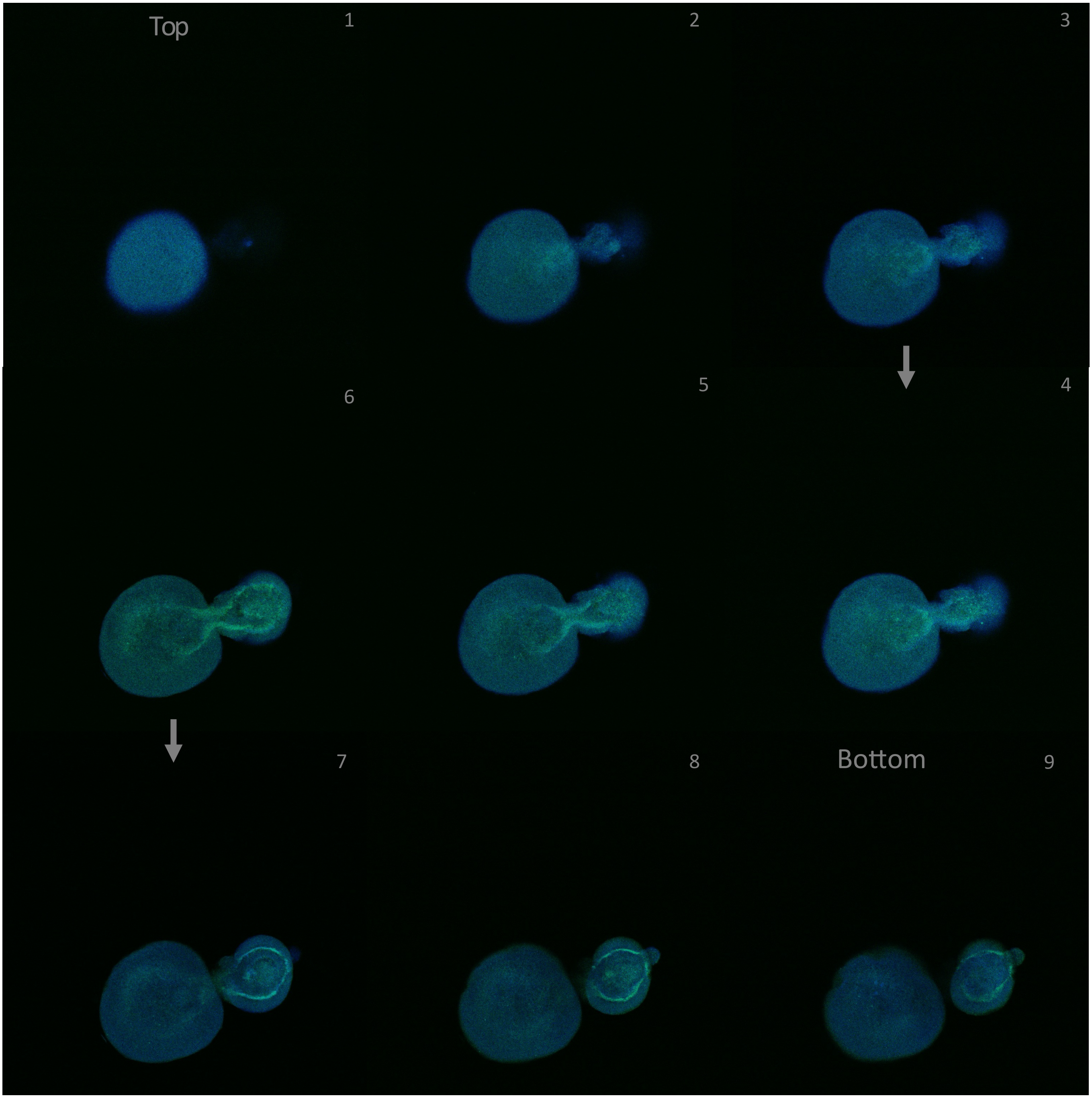


**Extended Data Fig. 3 | Confocal imaging of a conjoined pair of fully formed Embryoid Bodies (EBs)** with an genome integrated Cardiomyocyte enhancer within a base TRANC-ER construct (intron + polyA) in ChrX.

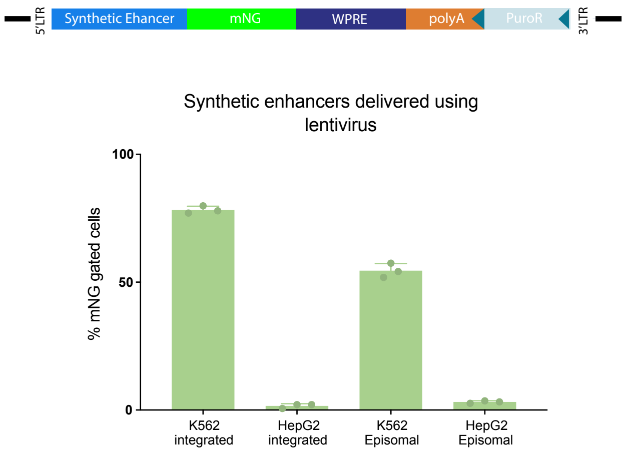


**Extended Data Fig, 4 | Lentiviral delivery of synthetic TRANCER constructs.** (Top) Diagram of the lentivirus TRANCER construct used in this experiment. (Bottom) Demonstration of TRANCER delivery using top-performing synthetic enhancers [51]. This panel shows that TRANCER constructs can be efficiently delivered and expressed in K562 cells via lentiviral transduction. Both integrated and episomal versions of the lentiviral TRANCER exhibit measurable reporter activity, confirming that lentiviral systems provide a viable platform for TRANCER delivery and functional expression. Data represent n = 3 biological replicates; error bars indicate SEM.

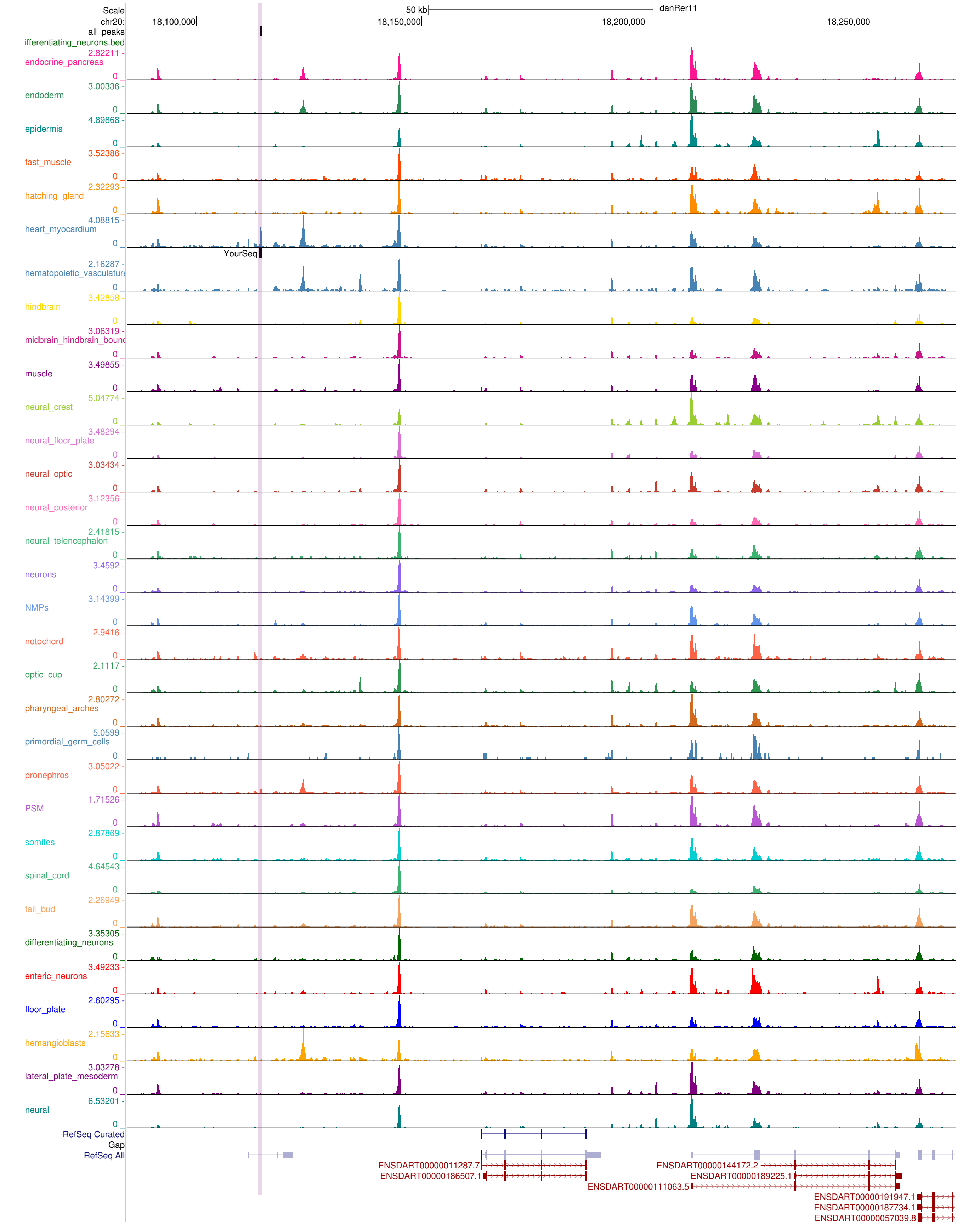


**Extended Data Fig. 5 | UCSC genome tracks from the Zebrahub 10x multiome resource.** The scATAC-seq data were re-analysed and visualised. The top enhancer region (chr20:18,114,062–18,114,563), highlighted in pink, was cloned into a TRANCER construct and used for subsequent experiments.

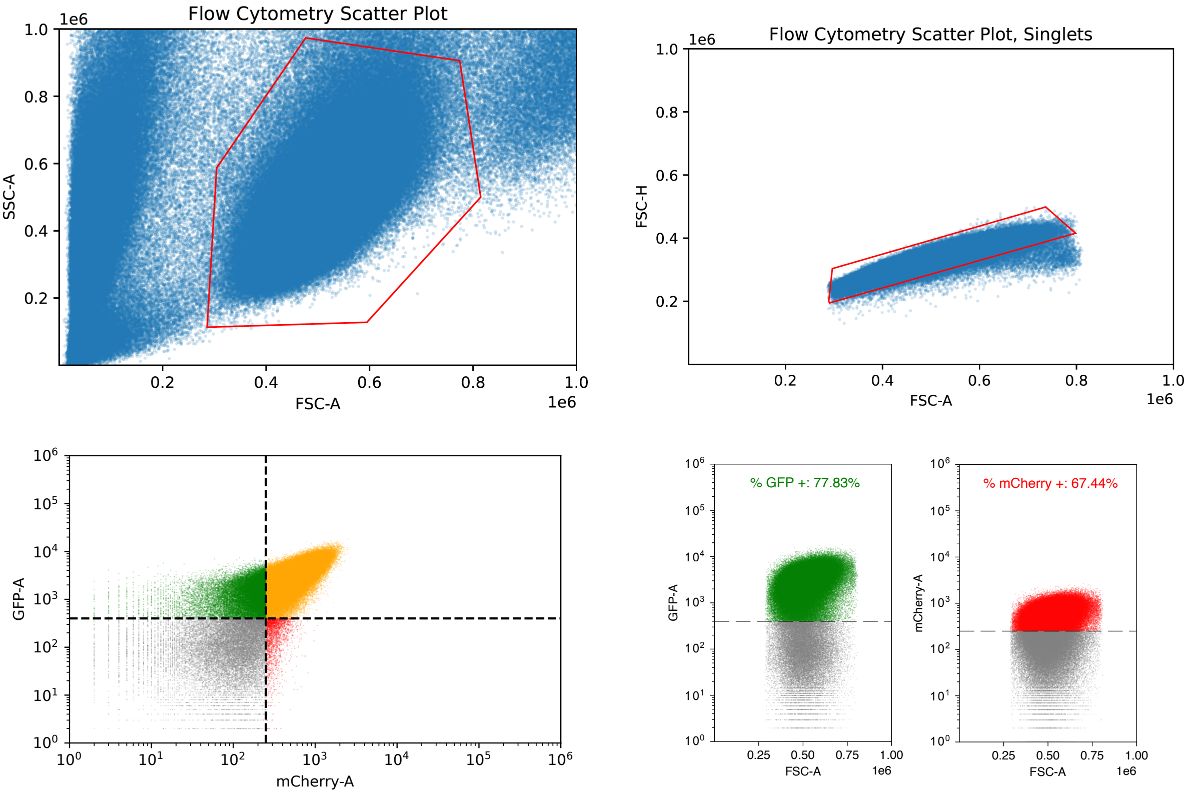


**Supplementary Figure 1 | Flow cytometry approach.** Flow cytometry was performed using an Attune NxT flow cytometer (Thermo Fisher Scientific). Cells were resuspended in PBS supplemented with fetal bovine serum (1:1) and kept on ice prior to acquisition. Data were analyzed using the FlowCytometryTools Python package.

For all experiments, debris was excluded based on forward and side scatter (FSC/SSC) parameters, followed by singlet discrimination using FSC-A versus FSC-H gating.

**Fluorescent reporter analysis:** Cells expressing mNeonGreen (mNG) and/or mCherry were analysed following live-cell and singlet gating. Fluorescence-positive populations were defined using negative controls. For simultaneous detection of both reporters, bivariate plots were generated with mCherry on the x-axis and mNG on the y-axis, and quadrant gates were used to identify single-positive, double-positive, and double-negative populations. Compensation was not required.

**Surface staining for CD19:** For surface marker analysis, cells were stained with an APC-conjugated anti-CD19 antibody according to the manufacturer's instructions. Following live-cell and singlet gating, CD19-positive cells were identified in the APC channel. As APC fluorescence was detected in the same channel as mCherry, identical gating strategies and appropriate controls were used to define positive populations.

**T-cell killing assays:** For T-cell killing assays, T cells were pre-labeled with CellTrace Yellow prior to co-culture with target K562 cells. After co-incubation, cells were stained with Live/Dead Violet to assess viability. During analysis, T-cells cells were identified based on CellTrace Yellow fluorescence, allowing separation from unlabeled K562 cells. Live and dead K562 cells were then quantified based on Live/Dead Violet staining.
